# Fob1-dependent condensin recruitment and loop extrusion on yeast chromosome III

**DOI:** 10.1101/2022.05.12.491704

**Authors:** Manikarna Dinda, Ryan D. Fine, Shekhar Saha, Zhenjia Wang, Chongzhi Zang, Mingguang Li, Jeffrey S. Smith

## Abstract

Despite recent advances in single-molecule and structural analysis of condensin activity *in vitro*, mechanisms of functional condensin loading and loop extrusion that lead to specific chromosomal organization remain unclear. In *Saccharomyces cerevisiae*, the most prominent condensin loading site is the rDNA locus on chromosome XII, but its repetitiveness deters rigorous analysis of individual genes. An equally prominent non-rDNA condensin site is located on chromosome III (chrIII). It lies in the promoter of a putative non-coding RNA gene called *RDT1*, which is in a segment of the recombination enhancer (RE) that dictates *MAT*a-specific chrIII organization. Here, we unexpectedly find that condensin is recruited to *RDT1* through interactions with Fob1, Tof2, and cohibin (Lrs4/Csm1), a set of nucleolar factors that also recruit condensin to the rDNA. Using Micro-C XL, we uncover evidence for condensin-driven loop extrusion anchored by Fob1 and cohibin at *RDT1* that extends toward *MAT*a on the right arm of chrIII, supporting donor preference during mating-type switching. *S. cerevisiae* chrIII therefore provides a new platform for the study of programmed condensin-mediated chromosome conformation.

## Introduction

Condensins are highly conserved multi-subunit complexes that function in chromosome organization and segregation. In budding yeast, *Saccharomyces cerevisiae*, the single condensin complex consists of two V-shaped <u>S</u>tructural <u>M</u>aintenance of <u>C</u>hromosome (SMC) ATPase subunits (Smc2 and Smc4), and three regulatory non-SMC subunits (Ycs4, Ycg1 and Brn1). Brn1 serves as a kleisin subunit that bridges the ATP-binding domains of Smc2 and Smc4 (Lavoie et al., 2000). As with other SMC complexes, yeast condensin can form loops through an ATP-dependent loop-extrusion activity (Ganji et al., 2018). Condensin has also been hypothesized to self-associate to bring distant chromatin-bound loci into contact (Yatskevich et al., 2019).

A general mechanism for functional condensin loading onto chromatin that establishes specific chromosome conformation remains elusive, especially for vertebrates that require extensive condensation of their large genomes, perhaps making chromatin recruitment relatively non-specific. Condensin complexes in *S. cerevisiae* and *S. pombe*, which have much smaller genomes, are attracted to transcriptionally active chromatin regions through interactions with transcription factors and enhanced by nucleosome displacement (Iwasaki et al., 2015; Toselli-Mollereau et al., 2016). At tRNA genes in *S. cerevisiae*, for example, condensin association with RNA Pol III transcription factors and the Scc2/Scc4 cohesin loading complex mediates their clustering into several foci surrounding the nucleolus (D’Ambrosio et al., 2008; Haeusler et al., 2008). At the rDNA locus, located on the right arm of chromosome XII (chrXII), condensin is recruited to the intergenic spacer (IGS1/NTS1) through association with a nucleolar protein complex established by the DNA replication fork block protein Fob1 (Johzuka and Horiuchi, 2009). Fob1 binds to cis-acting DNA sequences (TER1 and TER2) at a strong DNA replication fork block (RFB) site where DNA double-strand breaks (DSB) can occur (Kobayashi and Horiuchi, 1996). Repair of the DSBs through homologous recombination produces extrachromosomal rDNA circles (ERCs) that accumulate in mother cells and negatively impact replicative lifespan (Defossez et al., 1999; Johzuka and Horiuchi, 2002; Sinclair and Guarente, 1997). Fob1 and its associated factors, cohibin (Lrs4, Csm1), and Tof2, also recruit cohesin and the nucleolar Sir2 histone deacetylase (HDAC) complex, RENT, to IGS1 in the rDNA (Huang et al., 2006), where they mediate transcriptional silencing of intergenic non-coding RNAs (Kobayashi and Ganley, 2005; Li et al., 2006), and exit from mitosis through the FEAR network (Shou et al., 1999; Straight et al., 1999). Like condensin, the cohibin complex forms an extended coiled-coil structure, but it does not have ATPase activity (Corbett and Harrison, 2012).

A third major condensin recruitment site in *S. cerevisiae* is the recombination enhancer (RE) located adjacent to *HML* on the left arm of chromosome III (chrIII) (Li et al., 2019; Wu and Haber, 1996). The other silent mating-type locus, *HMR*, is located on the right arm of chrIII. In haploid *MAT*a cells only (Li *et al.*, 2019), condensin binds to a segment of the RE (the right half) that increases the frequency of *HML* contacting the right arm of chrIII, including the *MAT*a locus (Belton et al., 2015; Li *et al.*, 2019). While not required (Szeto et al., 1997; Wu and Haber, 1996), this segment of the RE and its resulting structural contribution instead augment donor preference of mating-type switching in *MAT*a cells (Li *et al.*, 2019), whereby *HML* is primarily utilized as the homologous donor template in repairing the DNA double-strand break induced by HO endonuclease at *MAT*a, thus converting it to *MAT*α (Haber, 2012). Once cells are switched to *MAT*α, condensin association with the RE is permanently lost (Li *et al.*, 2019), consistent with the extreme chrIII conformational differences between *MAT*a and *MAT*α cells.

Condensin recruitment as well as Sir2 at the RE occurs within the promoter of a small *MAT*a-specific gene of unknown function called *RDT1* (Li *et al.*, 2019; Wilson and Masel, 2011). Induction of mating-type switching by HO expression correlates with loss of condensin and Sir2 enrichment from *RDT1* and elevated *RDT1* expression (Li *et al.*, 2019), suggesting the region acts as a locus control region (LCR) that coordinates transcriptional regulation with chromosomal architecture.

*MAT*a-specific enrichment of condensin at *RDT1* is dependent on a consensus binding site for the MADS-box transcription factor Mcm1 (Li *et al.*, 2019). MADS-box transcription factors generally partner with other proteins to regulate transcription. For example, Mcm1 activates *MAT*a-specific genes in *MAT*a cells. In *MAT*α cells, however, Mcm1 forms a repressive heterodimer with the α2 protein (Mcm1/ α2) that inactivates the same *MAT*a-specific genes (Elble and Tye, 1991), including *RDT1* and another non-coding RNA gene (R2) the left half of the RE that regulates donor preference of mating-type switching (Ercan et al., 2005; Szeto *et al.*, 1997). Mcm1 also regulates cell cycle genes independent of mating type, including the cyclins *CLN3* and *CLB2* (Messenguy and Dubois, 2003). Since condensin and Sir2 were not enriched at such promoters in previous ChIP-seq datasets (Li *et al.*, 2019; Li et al., 2013), we hypothesized that additional factors partner with Mcm1 to recruit and load condensin at the *RDT1* promoter. Herein, we unexpectedly find that nucleolar factors cohibin, Fob1, and Tof2, which recruit condensin to the rDNA locus, also directly recruit condensin to the *RDT1* promoter in *MAT*a cells to direct chrIII conformation and augment donor preference.

## Results

We previously described strong *MAT*a-specific condensin and Sir2 enrichment at the *RDT1* promoter that was dependent on an upstream Mcm1/α2 binding site (*DPS2*; Fig 1A). Deleting 100bp of sequence underlying the Sir2 and condensin ChIP-seq peaks (100bpΔ; chrXII coordinates 30701-30800), while leaving the Mcm1/α2 site intact, also significantly disrupted condensin and Sir2 recruitment (Li *et al.*, 2019), suggesting one or more unknown factors cooperate with Mcm1 (Fig 1A). We initially focused on a putative Mcm1-Lrs4 interaction identified through a TAP-tag pulldown screen (Chan et al., 2011). Lrs4 and Csm1 are coiled-coil proteins that form a v-shaped complex known as cohibin in mitotic cells (Fig 1B; (Corbett and Harrison, 2012; Mekhail et al., 2008)). Lrs4 and Csm1 were of interest because they help recruit condensin and cohesin to the rDNA through interaction with Fob1 (Huang *et al.*, 2006; Johzuka and Horiuchi, 2009). Cohibin also physically interacts with Sir2 through the RENT and SIR complexes at the rDNA and telomeres, respectively (Chan *et al.*, 2011; Huang *et al.*, 2006). We confirmed the putative Mcm1-cohibin interaction with co-immunoprecipitations between Myc-tagged Mcm1 and Flag-tagged Csm1 (Fig 1C). Myc-tagged Lrs4 and Csm1 were also highly enriched at the *RDT1* promoter in quantitative ChIP assays, specifically in *MAT*a cells (Fig 1D). As a control, Lrs4-myc enrichment at the rDNA intergenic spacer was not mating-type specific (Fig 1E).

**Figure 1.**
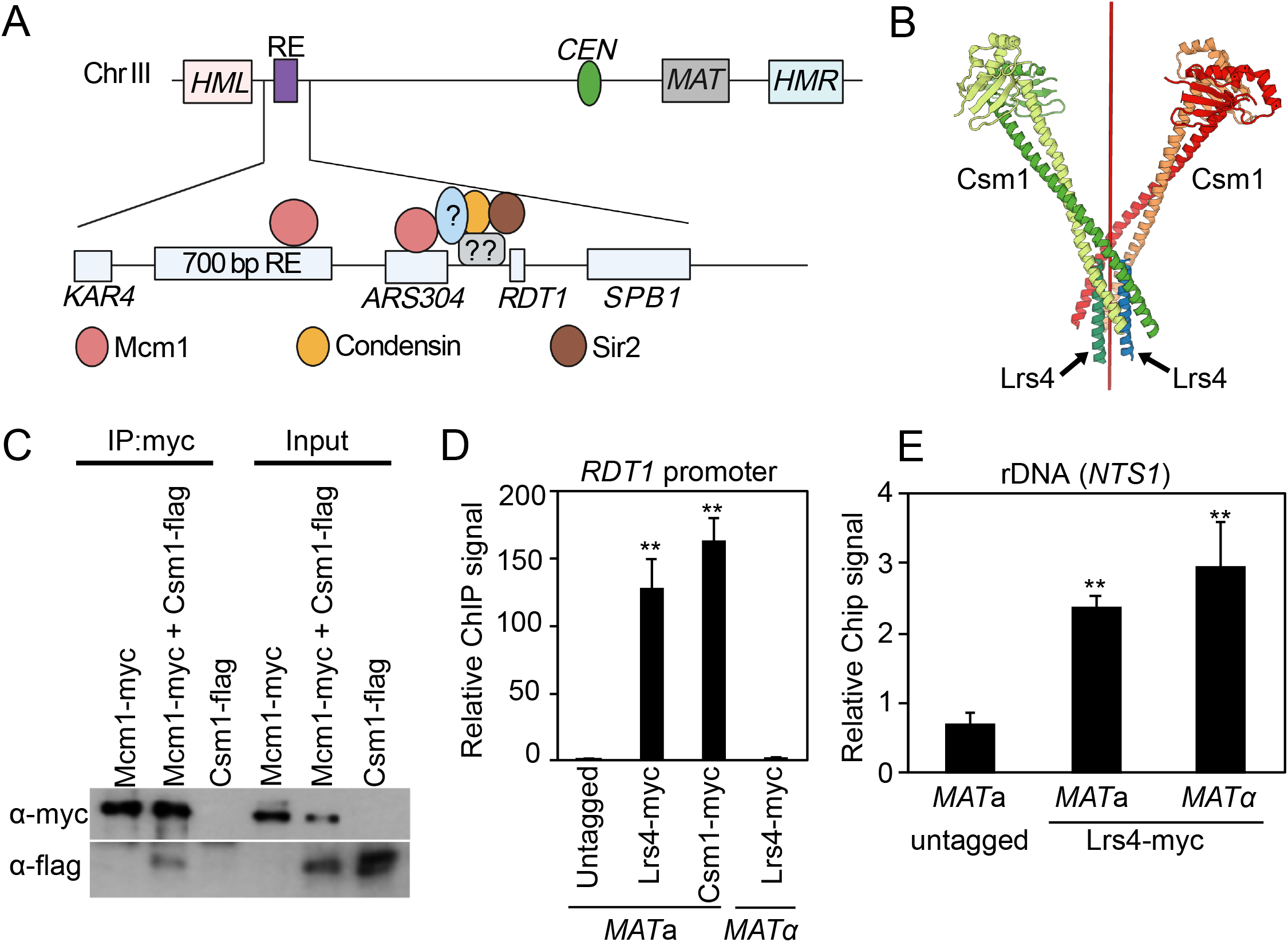
Cohibin binds to the *RDT1* promoter on chrIII specifically in *MAT*a cells. (*A*) Schematic of chrIII in *MAT*a cells. The recombination enhancer (RE) region lies between *KAR4* and *SPB1* consists of the minimal 700 bp element required for donor preference, two Mcm1/α2 binding sites, *ARS304*, and a Sir2/condensin site at the *RDT1* promoter. Uncharacterized factors are proposed to assist Sir2 and condensin recruitment by Mcm1. (*B*) X-ray crystallographic structure of cohibin consisting of 4 Csm1 subunits and 2 short Lrs4 subunit peptides. Red line indicates the axis of symmetry. (*C*) Immunoblot showing that Mcm1-myc coimmunoprecipitates with Csm1-flag. Loaded inputs are 5% of cell lysate used for the IP. (*D-E*) Enrichment of Lrs4-myc and Csm1-myc at the *RDT1* promoter or *NTS1* regions of the rDNA. (**p<0.005 relative to untagged strain).

### Cohibin is required for condensin enrichment at the *RDT1* promoter

The interaction of cohibin with Mcm1 and its strong enrichment at *RDT1* in *MAT*a cells supported a model whereby cohibin acts as a bridge between Mcm1 and the condensin/Sir2 complexes (Fig 1A). To investigate if cohibin was contributing to condensin recruitment specifically at *RDT1* or perhaps more generally across the genome, we performed ChIP-seq for a Myc-tagged subunit of condensin (Fig 2A; Brn1-13xmyc) in *MAT*a WT and *lrs4*Δ strains, as well as C-terminally tagged Lrs4 (Lrs4-13xmyc) strain. Myc-tagged centromeric protein, Ctf3, was used as a control for “hyper-ChIPable” sites that appear in yeast ChIP-seq datasets (Teytelman et al., 2013). We identified >700 sites that overlapped between Lrs4-myc and Brn1-myc after excluding Ctf3-myc peaks (Fig 2B, left Venn diagrams). A large proportion of the overlapping peaks showed differential Brn1-myc binding in the absence of Lrs4 (Fig 2B, right Venn diagrams). If cohibin was recruiting condensin genome-wide, then we expected Brn1-myc to decrease at most sites in the *lrs4*Δ mutant. However, only 7 Brn1-myc sites had a log2 fold change below −1 (Fig 2C), suggesting that cohibin-dependent recruitment of condensin is locus-specific. All the differential Brn1-myc sites are listed in Table S1.

**Figure 2.**
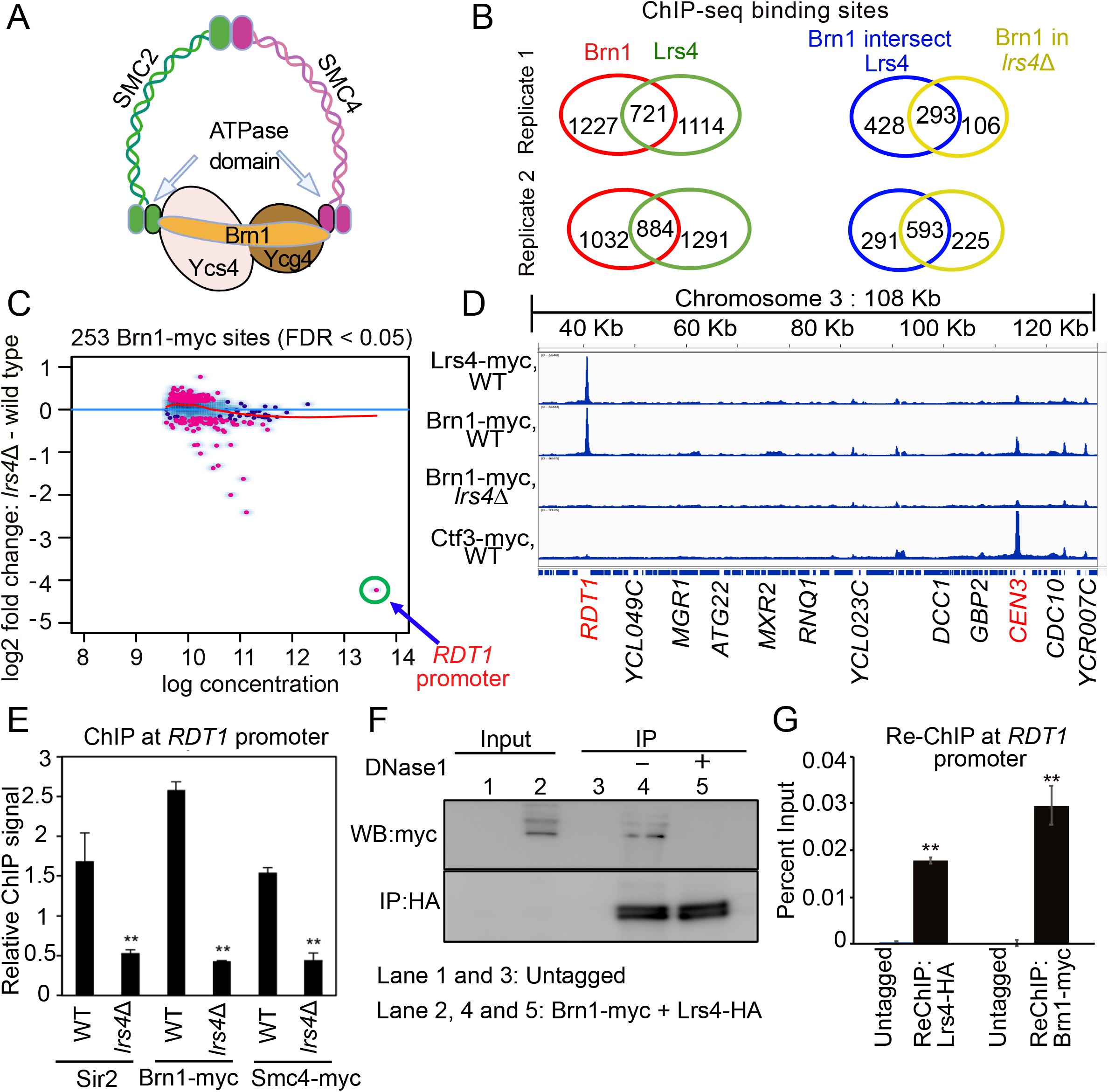
Cohibin recruits condensin to the *RDT1* promoter. (*A*) Schematic of budding yeast condensin subunits. (*B*) Venn diagrams of overlapping ChIP-seq peaks for Brn1-myc and Lrs4-myc from two independent replicates on the left. Among the overlapping peaks, the number of those that change in the *lrs4*Δ mutant is indicated in right Venn diagrams. Peaks that overlap with Ctf3-myc peaks were subtracted from each dataset. (*C*) MA plot indicating differential Brn1-myc binding between WT and *lrs4*Δ. Red dots are peaks significantly different between WT and *lrs4*Δ with FDR < 0.05. Blue dots are peaks not differentially bound. *RDT1* promoter peak is highlighted. (*D*) IGV screenshot of chrIII peaks for Brn1-myc, Lrs4-myc, Ctf3-myc, and Brn1-myc in the *lrs4*Δ mutant. Ctf3 is a centromere-specific control. (*E*) ChIP analysis at *RDT1* showing decreased enrichment of Sir2, Brn1-myc, and Smc4-myc in *lrs4*Δ mutant. (*F*) Co-IP assays from cell lysates were performed with an untagged strain or a strain with Lrs4-HA/Brn1-myc. DNase I digestion disrupts the coimmunoprecipitation. (*G*) Quantitative ChIP-reChIP PCR analysis of condensin (Brn1-myc) and cohibin (Lrs4-HA) co-occupancy at the *RDT1* promoter. (*p<0.05, **p<0.005).

Interestingly, the strongest Lrs4-dependent binding site was the *RDT1* promoter (Fig 2C). When displayed on IGV, overlapping distinctive peaks for Lrs4-myc and Brn1-myc were clearly observed at *RDT1* on chrIII, with smaller overlapping peaks at several other loci (Fig 2D). As expected from the MA plot in Fig 2C, the strong Brn1-myc peak at *RDT1* was eliminated in the *lrs4*Δ mutant (Fig 2D). A similar ChIP-seq pattern was observed for the rDNA at NTS1 (Fig S1A), where cohibin was known to recruit condensin (Johzuka and Horiuchi, 2009). We validated the loss of condensin at *RDT1* by quantitative ChIP-PCR for Brn1-myc and Smc4-myc subunits in the *lrs4*Δ mutant (Fig 2E) and observed similar depletion for endogenous Sir2 (Fig 2E). Enrichment of Lrs4 at the *RDT1* promoter was highly specific, as confirmed by a lack of detectable Lrs4-myc ChIP signal at the adjacent *RDT1* open reading frame (Fig S1B).

Importantly, Ctf3-myc was not enriched at *RDT1*, but as expected, was strongly bound to the chrIII centromere (Fig 2D; *CEN3*). Ctf3-myc also produced non-specific peaks that overlapped with Lrs4-myc and Brn1-myc (Fig 2D), likely examples of “hyper-ChIPable” loci. Cohibin co-localizes with condensin at mitotic and meiotic kinetochores in *S. cerevisiae* and *S. pombe* where they promote proper chromosome segregation (Brito et al., 2010; Peplowska et al., 2014). However, only in *S. pombe* was cohibin (Pcs1/Mde4) directly implicated in recruiting condensin to kinetochores (Tada et al., 2011). Here, we observed Lrs4-myc and Brn1-myc enrichment at most centromeres, including *CEN3* (Fig 2D). Moreover, the Brn1-myc peak at *CEN3* was clearly diminished in the *lrs4*Δ mutant (Fig 2D), suggesting that cohibin may also participate in condensin recruitment to centromeres in *S. cerevisiae*, though these sites were discarded from Fig 2B and C because they are also Ctf3 binding sites. To confirm that cohibin and condensin co-associate on chromatin, as predicted for a recruitment/loading function, we first performed a co-immunoprecipitation experiment showing that the interaction between Brn1-13xmyc and Lrs4-4xHA was eliminated by DNaseI treatment (Fig 2F). Second, a ChIP-reChIP assay demonstrated their co-occupancy on the *RDT1* promoter (Fig 2G).

### Cohibin promotes mating-type switching

We previously showed that acute depletion of Brn1 fused to an auxin-inducible degron (AID) slowed the rate of mating-type switching and moderately impaired donor preference (Li *et al.*, 2019). Since Lrs4 was required for condensin recruitment at chrIII (Fig 2), we predicted a similar switching-impaired phenotype should be observed with *lrs4*Δ. As shown in Fig S2A, conversion from *MAT*a to *MAT*α was slowed in the *lrs4*Δ mutant, but not as severely as *sir2Δ*, which strongly inhibits switching due to HO endonuclease cutting at the *HML* and *HMR* donor templates. Donor preference for switching to *MAT*α was also partially impaired by *lrs4*Δ, as compared to the full defect when the RE is deleted (Fig S2B and C). Deleting the *RDT1* promoter region (100bpΔ) moderately impaired donor preference without slowing the switching process (Li *et al.*, 2019), indicating the two phenotypes are separable. Therefore, we hypothesize the direct consequence of preventing cohibin-dependent condensin recruitment to *RDT1* is perturbed donor preference, whereas the switching delay is likely due to pleiotropic effects.

### Fob1 recruits a nucleolar condensin loading complex to chromosome III

If interaction between Mcm1 and cohibin was the sole mechanism of bringing condensin to *RDT1*, then we would have expected Lrs4-myc enrichment at other known Mcm1 sites, including an Mcm1 site (DPS1) within the left half of the RE. This was not the case (Fig 2D). Instead, our model for condensin recruitment to *RDT1* allowed for contributions from one or more additional factors, including other site-specific DNA binding proteins (Fig 1A). Condensin and cohibin are both targeted to the RFB site in NTS1 by hierarchical interactions with Fob1 and Tof2, with Fob1 providing the site-specific DNA binding activity (Fig 3A, left panel; (Huang and Moazed, 2003; Johzuka and Horiuchi, 2009)). We therefore hypothesized that a similar hierarchy of Fob1-dependent recruitment occurred at *RDT1*, with mating-type specificity added by Mcm1 (Fig 3A, right panel). Fob1 had no previously known function outside the nucleolus/rDNA, but in ChIP assays we observed exceptionally strong *MAT*a-specific enrichment of Fob1-13xmyc at *RDT1*, compared to mating-type independent enrichment at NTS1 (Fig 3B). ChIP was also used to test whether a similar hierarchy of Fob1-dependent condensin recruitment was occurring at *RDT1* (Fig 3C) and rDNA (Fig 3D). Deleting *FOB1* or *TOF2* eliminated Lrs4-myc and Brn1-myc enrichment at *RDT1*, and a similar pattern was observed at the rDNA (NTS1), though the Lrs4-myc and Brn1-myc depletion was not absolute, perhaps because rDNA ChIP assays average the PCR signal from ∼150 to 200 repeats, or additional factors are involved in condensin and cohibin recruitment to the rDNA.

**Figure 3.**
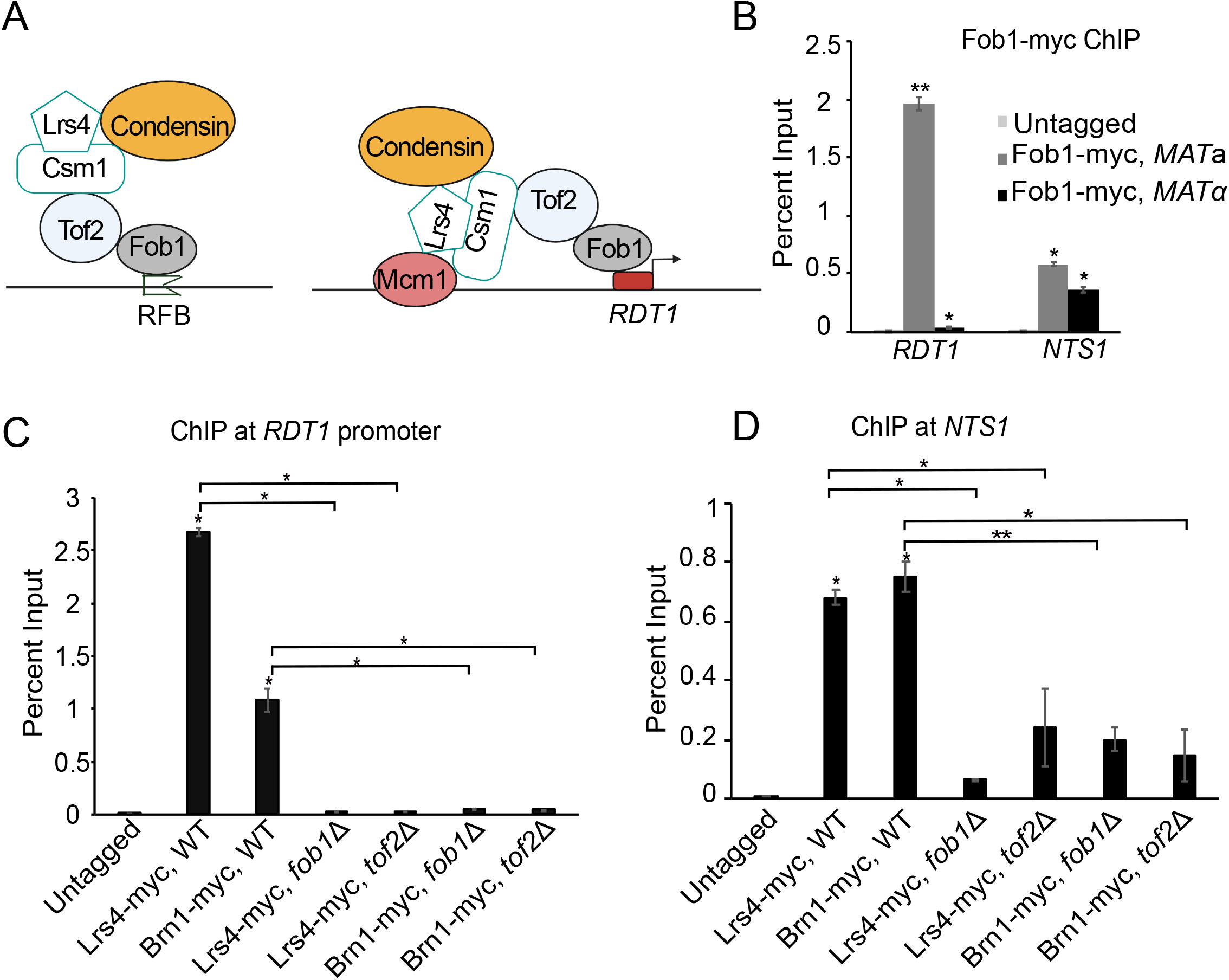
Fob1 recruits cohibin and condensin to the *RDT1* promoter. (*A*) Hierarchical models of condensin recruitment to the rDNA and *RDT1* promoter. The crystal structure of cohibin is depicted to emphasize its ability to bridge Mcm1 with other factors in the complex. (*B*) ChIP of Fob1-myc at *RDT1* promoter and *NTS1* in *MAT*a and *MAT*α cells. (*C-D*) ChIP analysis of Lrs4-myc and Brn1-myc enrichment at the *RDT1* promoter or *NTS1* showing dependence on *FOB1* and *TOF2*. (*p<0.05, **p<0.005).

### Fob1 directly binds to *RDT1* promoter sequence

Fob1 binding to rDNA occurs at the RFB site in NTS1, consisting of 2 replication fork termination sites (Fig 4A; TER1 and TER2) (Ward et al., 2000). These sites do not match sequence within the *RDT1* promoter, and due to TER1 and TER2 being the only previously known Fob1 binding sites, there is no consensus binding sequence for this protein. To test if Fob1 directly binds the *RDT1* promoter (Fig 4A, lower), we performed electrophoretic mobility shift assays (EMSA). GST-Fob1 was overexpressed in yeast and purified by affinity chromatography (Fig 4B), then incubated *in vitro* with small biotin-labeled PCR products derived from the *RDT1* promoter (chrIII coordinates 30712 to 30768) or RFB (chrXII coordinates 460533 to 460574) (Fig 5C). GST-Fob1 produced two band shifts with the RFB and *RDT1* probes, though they were qualitatively different (Fig 4D). The top RFB band shift was competed away with unlabeled RFB probe, while the weak faster migrating band appeared non-specific. Both band shifts with the longer *RDT1* probe were competed away by unlabeled *RDT1* probe, suggesting the presence of two specific binding sites, one stronger than the other. Importantly, purified GST control protein did not bind to either probe. In summary, this result shows that Fob1p directly binds to the *RDT1* promoter sequence, consistent with the hierarchical model for condensin recruitment defined by ChIP (Fig 3A).

**Figure 4.**
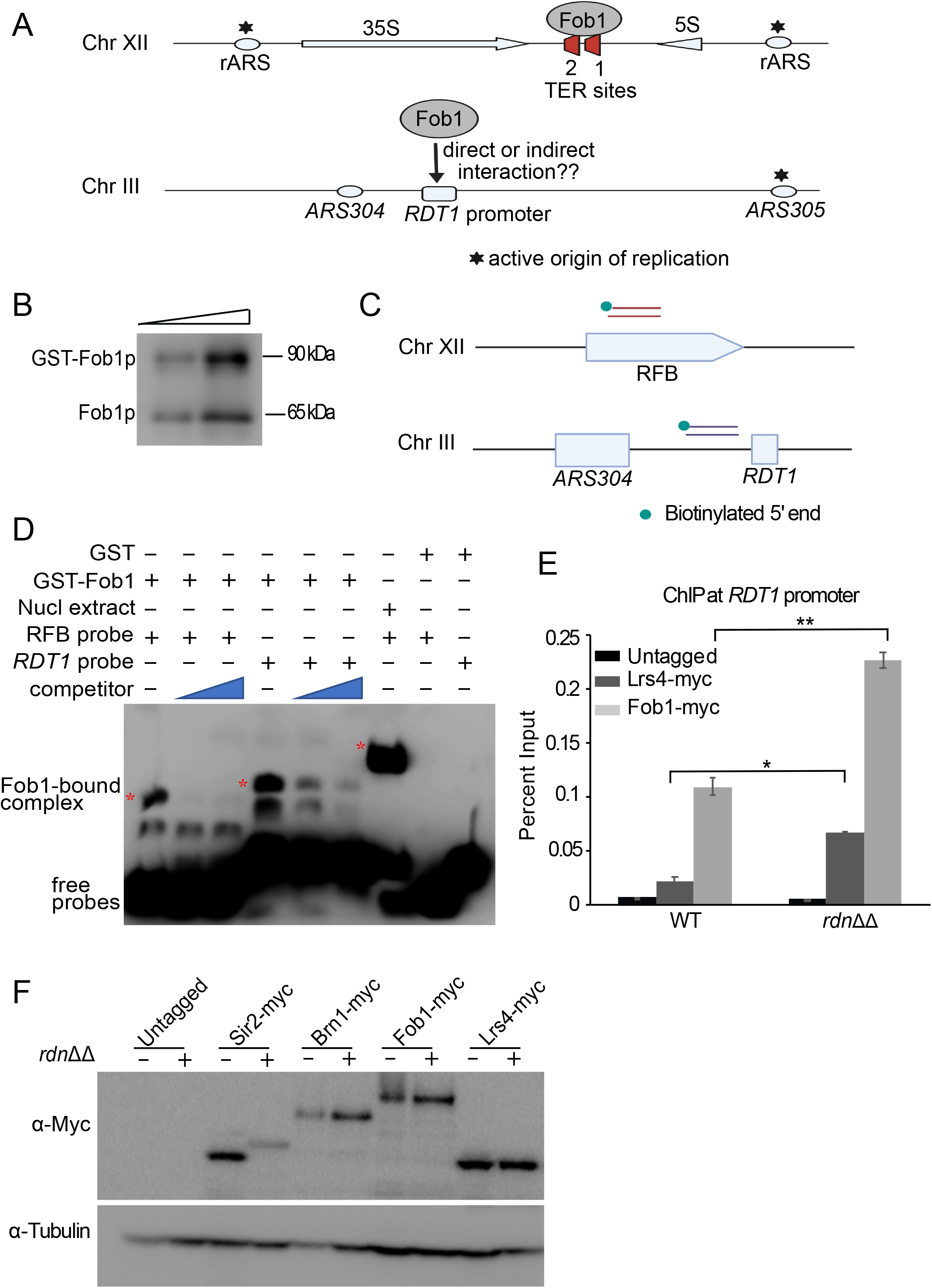
Recombinant Fob1 directly binds to the *RDT1* promoter sequence *in vitro*. (*A*) Schematic of Fob1 binding to TER1 and TER2 sites of the replication fork block (RFB), and the *RDT1* promoter. (*B*) Western blot of yeast-purified GST-Fob1p using α-Fob1 antibody. (*C*) Schematic for the design of biotin labelled probes used for EMSA. (*D*) EMSA with biotin-labeled probes and purified GST-Fob1p, GST, or nuclear extract control. Competitor probes were unlabeled. The *RDT1* probe is longer than RFB. (*E*) ChIP of Lrs4-myc and Fob1-myc at the *RDT1* promoter in a strain (NOY891) lacking the rDNA array (*rdnΔΔ*) or W303 WT control. ChIP signal is relative to the input chromatin signal expressed as percent input (*p<0.05, **p<0.005). (*F*) Western blot showing steady state Brn1-myc, Lrs4-myc, Fob1-myc, and Sir2-myc levels in WT and *rdnΔΔ* strains. α-tubulin is used as a loading control.

**Figure 5.**
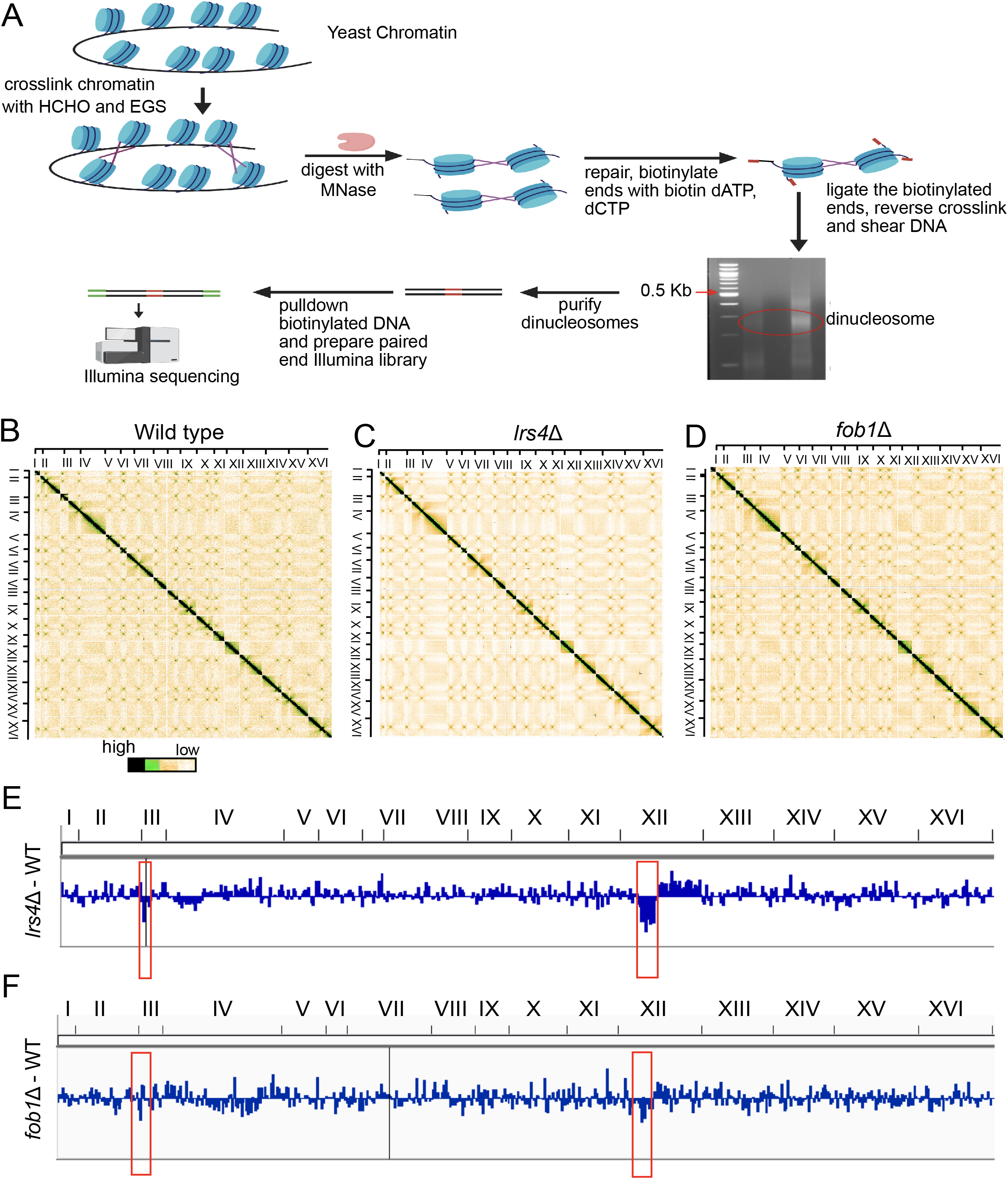
Micro-C XL analysis of the yeast genome in *lrs4*Δ and *fob1*Δ mutants. (*A*) Overview of the Micro-C XL method. (*B-D*) Genome-wide interaction heatmaps of Micro-C XL data in exponentially growing wild type, *lrs4*Δ and *fob1*Δ cells. Each pixel represents normalized chromatin contacts between 10 kb bins. (*E-F*) Differential Chromatin Interaction (DCI) scores were calculated for normalized contact frequencies of 5kb target across +/-200kb flanking regions, comparing *lrs4*Δ or *fob1*Δ mutants with WT. The negative DCI scores observed in chromosome III and chromosome XII are boxed in red.

We next tested if *RDT1* was in competition with the repetitive rDNA for the predominantly nucleolar condensin recruitment complex. Fob1 and Lrs4 were C-terminally tagged with 13xMyc in a strain lacking the chromosomal rDNA array (*rdnΔ*Δ) and kept viable by a plasmid expressing 35S rRNA precursor from a galactose-inducible RNA Pol II promoter (Wai et al., 2000). ChIP assays for *RDT1* promoter enrichment in the WT and *rdnΔ*Δ strains were then performed (Fig 4E). Interestingly, Fob1-myc and Lrs4-myc showed significantly improved enrichment at *RDT1* in the *rdnΔ*Δ mutant while the overall cellular protein levels were unaffected (Fig 5F and G). This contrasted with the Sir2-myc positive control, which showed reduced protein in the *rdnΔ*Δ mutant, as predicted for strains with low rDNA copy number (Iida and Kobayashi, 2019; Michel et al., 2005). From this experiment we concluded that the single copy *RDT1* promoter region competes the repetitive rDNA locus limiting Fob1 and cohibin, suggesting that rDNA copy number could potentially contribute to regulation of chromosome III structure.

### Cohibin and Fob1 dictate the conformation of chromosomes III and XII

Given the requirements for cohibin and Fob1 in condensin recruitment to the *RDT1* promoter on chrIII and the rDNA on chrXII, we hypothesized that deleting *LRS4* or *FOB1* would significantly disrupt the structure of both chromosomes. To investigate changes in 3D conformation, we utilized Micro-C XL (Fig 6A), which takes advantage of micrococcal nuclease digestion of crosslinked chromatin and purification of ligated dinucleosomes to enhance resolution (Hsieh et al., 2016). Normalized interaction frequencies within 10 kb bins across the entire genome were first visualized in heatmap plots for WT, *lrs4*Δ and *fob1*Δ strains. Interchromosomal interaction patterns were generally similar between the three strains (Fig 5B-D). For example, absence of Lrs4 or Fob1 did not reduce the number of centromere-centromere or telomere-telomere interactions among the top 10% of overall interactions (Fig S3A and B), indicating gross interchromosomal organization remained intact. We also quantified differential intrachromosomal contacts between WT and the *lrs4*Δ or *fob1*Δ mutants using BART3D (Wang et al., 2021). A differential chromatin interaction (DCI) score was calculated for each 5kb segment across the genome to measure the difference in normalized contact between each segment and its +/-200kb flanking regions (Fig 5E-F). Segments with the strongest negative DCI scores, representing the most decreased interactions in the *lrs4*Δ mutant compared to WT (Fig S3C), were predominantly from a region on the right arm of chrXII between the centromere and rDNA array (Fig 6E and S3E), corresponding to a large condensin-dependent chromatin loop (Fine et al., 2019; Lazar-Stefanita et al., 2017; Wang *et al.*, 2021). Furthermore, the *RDT1*-containing chrIII segment (coordinates 30000 to 35000) showed the 2^nd^ strongest negative DCI score (Fig 5E and S3E). Though not as striking due to a larger number of segments with negative DCI scores (Fig S3D), chrIII and XII were still among the most affected in the *fob1*Δ mutant (Fig 5F and S3F), with the RE-containing segment the top hit (Fig 5F and S3F). Taken together, we conclude that Lrs4 and Fob1 both dictate the structures of chrIII and XII, most likely through localized recruitment and loading of condensin.

**Figure 6.**
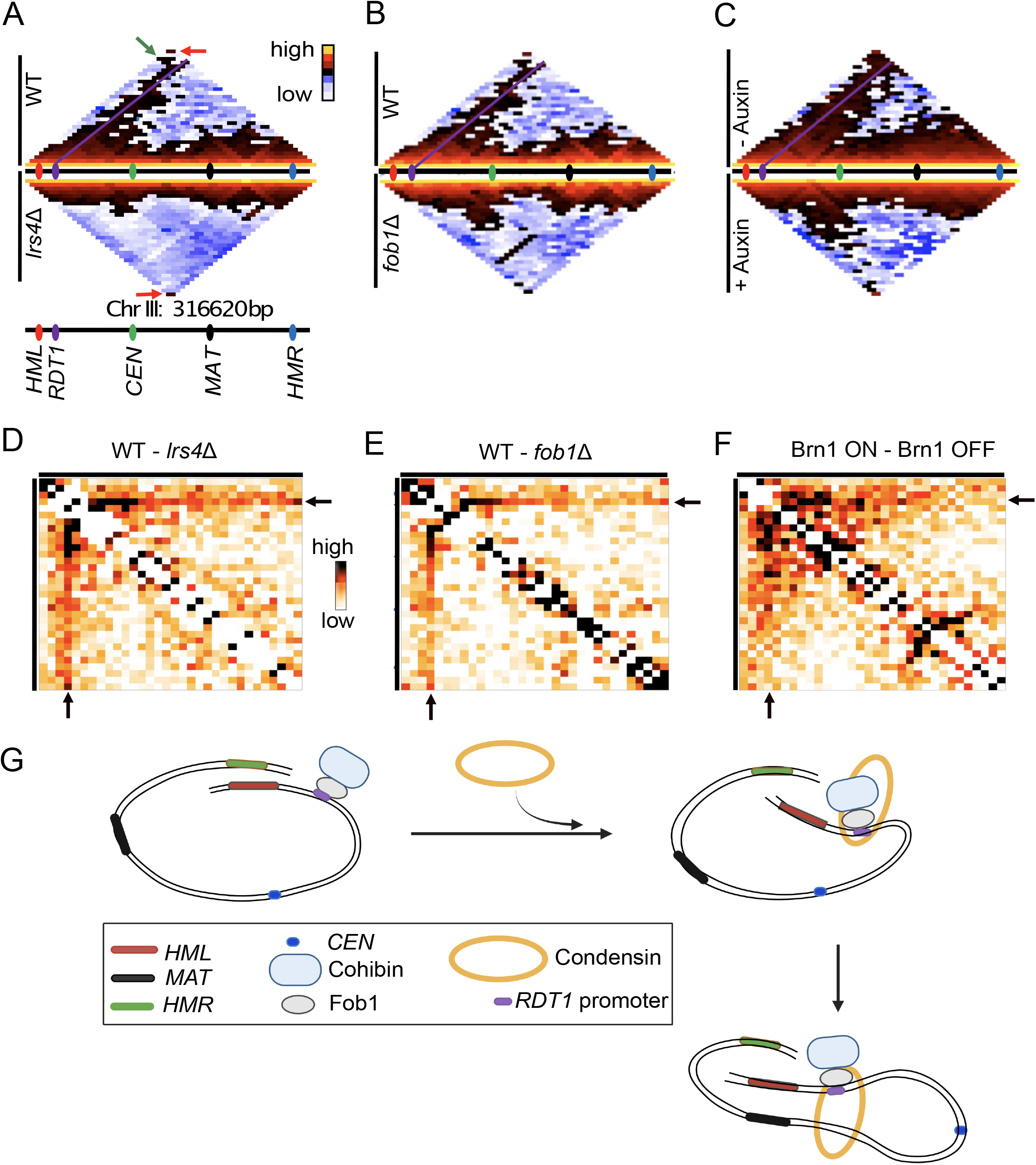
Lrs4, Fob1, and condensin control chromosome III conformation in *MAT*a cells. (*A*) Interaction heat map for chromosome III at 10 kb resolution comparing WT (top) and *lrs4*Δ (bottom) strains. In the WT map, a green box indicates the *HML-HMR* interaction, a purple line indicates interaction of the *RDT1*-containing bin with the rest of chrIII. Red boxes indicate the telomere-telomere interaction. (*B*) ChrIII interaction heat maps for WT and *fob1*Δ strains. (*C*) Interaction heat maps for chrIII during mating-type switching after depleting Brn1 using the auxin-inducible degron (AID) system. (*D-E*) Subtraction of *lrs4*Δ or *fob1*Δ interaction frequencies from WT reveals a rightward gradient of interaction anchored at the *RDT1* bin. (*F*) Subtraction of the Brn1-AID depletion interaction frequencies from non-depleted control reveals strong reorganization across the left arm of chrIII. Black indicates higher interactions in WT compared to the mutants. (*G*) Loop extrusion model for condensin recruited to the *RDT1* promoter by Fob1 and cohibin.

Previous 3C and Hi-C analyses of chrIII conformation in *MAT*a and *MAT*α cells identified a prominent interaction between the subtelomeric *HML* and *HMR* loci (Belton *et al.*, 2015; Miele et al., 2009). Deleting the *RDT1* promoter (100bpΔ mutant) was sufficient to disrupt the HML-HMR interaction at least partially in *MAT*a cells, freeing *HMR* to interact with the *MAT*a locus (Li *et al.*, 2019). This mutant also disrupted contacts between *HML* and the right arm of chrIII, including the *MAT*a locus (Li *et al.*, 2019). To investigate if Fob1 and Lrs4 were required for these specific structures, iteratively corrected chrIII Micro-C XL contact maps (10kb bins) for each mutant were directly compared to WT (Fig 6A and B). With the higher resolution of Micro-C XL, the WT pattern now delineated an apparently rightward-extended path of interaction anchored at the RE-containing bin, as well as the expected *HML*-*HMR* and telomere-telomere interacting bins, which were offset from the line of RE-anchored interactions. Importantly, the *HML*-*HMR* and RE-anchored interactions were disrupted in the *lrs4*Δ and *fob1*Δ mutants (Fig 6A and B), but the telomere-telomere interaction was retained (Fig 6A and B). Subtraction plots of the Micro-C XL data revealed a gradient along the RE-anchored interaction path (Fig 6D and E), consistent with Lrs4-or Fob1-recruited condensin at the RE mediating loop extrusion (Fig 6G). To confirm that loss of condensin recruitment directly contributed to the altered chrIII structure, we performed Micro-C XL with a strain containing Brn1 C-terminally tagged with an auxin-inducible degron (AID). Following 30 minutes of Brn1-AID depletion during induction of mating-type switching (Fig 7B), a similar loss of RE-anchored interaction across chrIII was observed, though the condensin-dependent left arm contacts were expanded beyond the RE anchor bin, suggesting that condensin contributes more broadly to chrIII reorganization during the switching process.

**Figure 7.**
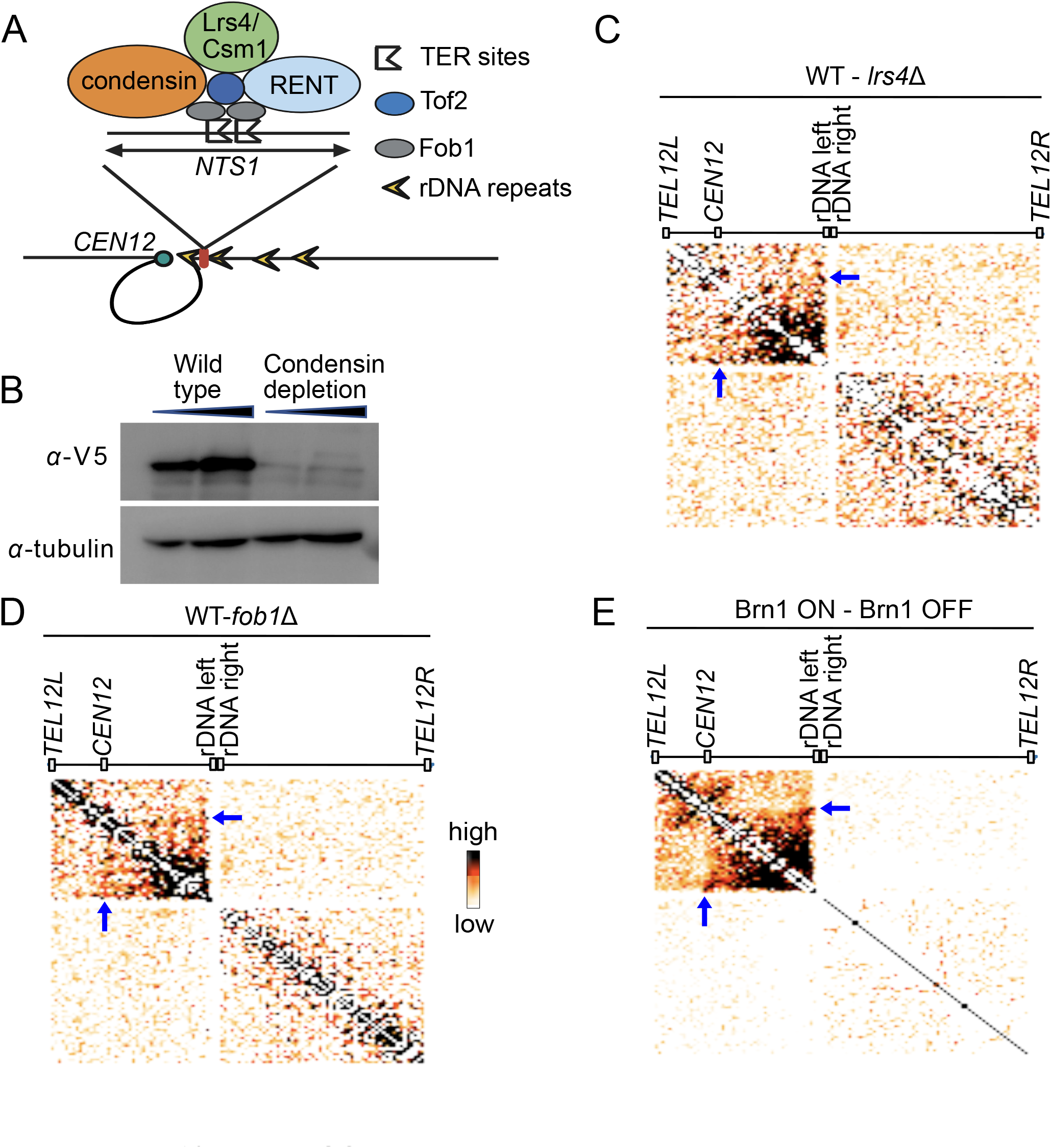
Micro-C XL analysis of chrXII conformation. (*A*) Schematic of condensin, cohibin, Tof2, and RENT recruitment to *NTS1* by Fob1 in the rDNA array. A putative TAD element or large loop between *CEN12* and the unique sequence flanking the rDNA on chrXII is also depicted. (*B*) Western blot showing condensin depletion after 30 min with α-V5 antibody that recognizes the AID fusion on Brn1. (*C-E*) Micro-C XL contact maps for chrXII at 10 kb resolution showing *lrs4*Δ, *fob1*Δ and Brn1-AID interaction frequencies subtracted from wild-type. Black indicates higher interactions in WT compared to the mutants and white indicate no change.

Since Fob1 and cohibin also recruit condensin to the rDNA, and condensin is required to form a putative TAD between the array and the chrXII centromere (Fig 7A), we generated subtraction plots for the chrXII Micro-C XL data between WT and the *lrs4*Δ mutant (Fig 7C), the *fob1*Δ mutant (Fig 7D), and Brn1-AID depletion strain (Fig 7E). Loss of the *CEN12*-rDNA loop was especially clear with acute Brn1-AID depletion, with little change to the right of the rDNA array. Large changes within the TAD structure were also observed with *lrs4*Δ and *fob1*Δ, but these mutants also caused diffuse alterations along the right arm and even between the two arms that could be due to secondary structural impacts beyond simply recruiting condensin, perhaps through their additional roles in recruiting cohesin to the rDNA (Huang *et al.*, 2006). Taken together, we conclude that, like the rDNA locus, Fob1 and cohibin recruit condensin to the *RDT1* promoter where it folds chrIII into a specialized conformation via loop extrusion.

## Discussion

Interphase chromosomes of multicellular organisms are organized into 3-dimensional domains ranging in size from kilobase scale chromatin loops to megabase scale topologically associated domains (TADs). Yeast chromosomes are also 3-dimensionally organized, but at a much smaller scale. Computational modeling, imaging, and Hi-C analyses predict a Rabl-like conformation whereby centromeres are clustered with the duplicated spindle pole bodies, and telomeres clustered into several perinuclear foci (Duan et al., 2010). Tethered centromeres limit spatial mobility of the chromosome arms (Verdaasdonk et al., 2013), which essentially fill limited space within the small yeast nucleus, resulting in chromosome “territories” that can be largely explained by geometrical constraints (Tjong et al., 2012). Despite such limitations, there are still multiple biochemically generated architectural features within the yeast genome, including abundant self-associating chromatin domains detected by Micro-C that only span across one to five genes and are dependent on chromatin remodeling factors (Hsieh et al., 2015; Hsieh *et al.*, 2016). There are also biochemically generated large-scale features, such as the rDNA array and the condensin-dependent TAD formed between the centromere and rDNA on chrXII (Fig 7; (Duan *et al.*, 2010; Fine *et al.*, 2019; Lazar-Stefanita *et al.*, 2017)). The *MAT*a-and *MAT*α-specific conformations of chrIII comprise the other major large-scale architectural features of the budding yeast genome (Belton *et al.*, 2015; Miele *et al.*, 2009).

### A shared mechanism of condensin recruitment between the rDNA and chrIII

Lrs4 (<u>L</u>oss of <u>r</u>DNA <u>s</u>ilencing 4) was first identified through a screen for genes that function in Sir2-dependent rDNA silencing (Smith et al., 1999). It was later found to be a subunit of the meiotic monopolin complex (Lrs4/Csm1/Mam1) responsible for monopolar attachment of microtubules to kinetochores at meiosis I (Rabitsch et al., 2003). We became interested in cohibin (Lrs4/Csm1) as a possible condensin and Sir2 recruitment factor for chrIII in *MAT*a cells because pull-down experiments with TAP-tagged Lrs4 and Csm1 subunits identified Mcm1 as a putative interacting protein (Chan *et al.*, 2011). Additionally, Lrs4, Csm1, and Tof2 were previously identified in a protein interaction network with Fob1 and the RENT complex (Huang *et al.*, 2006), and to be involved in both cohesin and condensin recruitment at NTS1 in the rDNA (Huang *et al.*, 2006; Johzuka and Horiuchi, 2009). While primarily localized at the rDNA/nucleolus, some of these factors have known functions outside the nucleolus. The Cdc14 subunit of RENT is partially redistributed outside the nucleolus during early anaphase by the FEAR network (D’Amours and Amon, 2004), and then further released at late anaphase by the Mitotic Exit Network (MEN) (Torres-Rosell et al., 2005). Similarly, cohibin is released from the nucleolus by the MEN network during anaphase, relocalizing to kinetochores to ensure faithful chromosome segregation (Brito *et al.*, 2010). Cohibin also interacts with telomeres where it promotes their clustering at the nuclear periphery during S and G1 through association with inner nuclear membrane proteins (Chan *et al.*, 2011). It remains to be determined if the RE/*RDT1* plays any role in the FEAR or MEN networks.

### Fob1 function at the RE

Recruitment of cohibin and condensin to the RE is the first non-nucleolar function described for Fob1. Cohibin recruitment to telomeres is dependent on the SIR proteins, not Fob1, and is enhanced in a *fob1*Δ mutant due to release of Sir2 from the rDNA (Chan *et al.*, 2011). There is also no evidence of cohibin recruiting condensin to telomeres, and we have not detected significant condensin enrichment at telomeric regions in ChIP-seq datasets. Therefore, the finding that Fob1 is required for cohibin and condensin recruitment at *RDT1* makes the underlying mechanism most like recruitment of these factors at rDNA. In this model, Fob1 provides the sequence-specific DNA binding function, either at the *RDT1* promoter or RFB TER sites at the rDNA. A key distinction is the Mcm1/α2 binding site upstream of *RDT1* likely controls the mating-type specificity. Assembly of an Mcm1/α2 heterodimer at the *RDT1* promoter in *MAT*α cells forms a repressive chromatin structure that either prevents DNA binding by Fob1 or blocks cohibin from interacting with Fob1 and Mcm1. The inability to load condensin therefore prevents the formation of *MAT*a-specific structural features. The absence of Mcm1/α2 regulation at the rDNA makes cohibin, condensin, and RENT recruitment independent of mating-type.

At the rDNA, Fob1 also acts as a unidirectional DNA replication fork block factor. Fob1 bound to the TER1/TER2 sites blocks forks moving in opposite direction of RNA Pol I transcription (Kobayashi and Horiuchi, 1996; Mohanty and Bastia, 2004). An autonomously replicating sequence (*ARS304*) is located immediately upstream of the Fob1 binding site at *RDT1* and is marked by the Mcm1/α2 binding site. Although bound by Mcm1, ARS304 and all other ARS elements to the left of *RDT1* are inactive (Vujcic et al., 1999), implying the region is replicated from the highly active *ARS305* located to the right of the RE. The RE or *RDT1* promoter region has not been implicated in fork blocking in WT strains, though pausing has been detected in a strain lacking Rrm3, a helicase that normally facilitates replication through non-histone protein complexes (Ivessa et al., 2003). Fork block activity of Fob1 at the rDNA is genetically separable from its role in silencing via RENT recruitment (Bairwa et al., 2010), so it is possible that both activities occur at *RDT1* under specific conditions such as a particular cell cycle stage or mating-type switching.

Fob1 and cohibin could potentially function in mediating chromatin organization of chrIII separate from their roles in recruiting condensin. For example, Fob1 was previously shown to mediate “chromosome kissing” between rDNA repeats through inter-repeat Fob1-Fob1 protein interactions (Choudhury et al., 2015). Cohibin is also thought to link different chromatin regions through protein-protein interactions mediated by globular domains of the Csm1 subunit, a mechanism by which cohibin-bound telomeres are clustered at the nuclear periphery through interaction with inner nuclear membrane proteins (Mekhail *et al.*, 2008). Given that *RDT1* is located only 17kb from *HML*, it is possible that the cohibin recruited to *RDT1* by Fob1 could potentially contribute similarly to the known localization of *HML* near the nuclear periphery (Bystricky et al., 2009).

### A model for condensin-mediated loop extrusion during mating-type switching

SMC complexes such as condensin and cohesin generate chromatin loops through ATP-dependent loop extrusion. Our Micro-C XL analysis of chrIII predicts that condensin anchored at the *RDT1* promoter performs loop extrusion in *MAT*a cells that extends toward the centromere and *MAT*a locus. While we could not perform a time course of increased DNA juxtaposition from the condensin-bound anchor point in the RE, as was previously done for the *Bacillus subtilis* SMC (condensin) complex loaded onto a specific ParB recruitment site on the circular chromosome (Wang et al., 2018), we did detect a discreet path of juxtaposition with rightward extension across a diminishing gradient. This path of juxtaposition was eliminated if condensing recruitment was blocked (*fob1*Δ or *lrs4*Δ) or if condensin was acutely depleted. Under such a model, loop extrusion anchored at the *RDT1* promoter would provide a mechanism that allows *HML* to scan across the right arm of chrIII, thus increasing the chance of contact with *MAT*a when mating-type switching is induced. In conclusion, we have established the *RDT1* promoter region of the RE as a new non-repetitive genetic system for studying programmed structural organization by condensin in yeast.

## Materials and methods

### Yeast strains and plasmids

Yeast strains listed in Table S2 were grown in yeast extract-peptone-dextrose (YPD) or appropriate synthetic complete (SC) dropout medium at 30°C. Gene deletions were constructed by PCR-mediated one-step replacement with *kanMX6*, *natMX6* or *hygMX6* cassettes. C-terminal epitope-tagging of endogenous genes was achieved by integrating 3xHA-*TRP*, 3xV5-*kanMX6*, or 13xMyc-*kanMX6* in place of the stop codon. To generate yeast strains expressing GST-Fob1, a *GST-FOB1* ORF segment was PCR amplified from pJSS92-8 (pGEX6P1 expressing GST-Fob1) and subcloned into *Hin*dIII/*Xba*I sites of pYES2.URA using the in-fusion method (Takara). The *rdn*ΔΔ yeast strain NOY891carrying pNOY353 was maintained on SC-trp 2% galactose (Wai *et al.*, 2000). Primers used in strain construction are listed in Table S3.

### Immunoblots and Immunoprecipitation

Samples for immunoblot analysis were prepared from whole cell extracts using the TCA/glass bead method as previously described (Li *et al.*, 2013). Immunoblots were performed with the following antibodies: anti-myc (Cell Signaling, #71D101:2,000), anti-HA (Cell Signaling, #C29F41:2000), M2 anti-FLAG (Sigma, #F1804, 1:2000), anti-Sir2 (Santa Cruz, 1:2000), or anti-tubulin (Millipore Sigma, #T5168 1:4000). Secondary antibodies were HRP-conjugated anti-mouse (Cell Signaling #7076S, 1:4000) or anti-rabbit HRP-conjugated (Cell Signaling #7074S, 1:4000). Protein–antibody conjugates were revealed by chemiluminescence (Immobilon Western Chemiluminescent HRP substrate, Millipore). To disrupt DNA-mediated protein interactions, cell lysates were digested with DNase 1 (100 μg/ml) at 37°C for 20 min prior to antibody incubation.

For immunoprecipitations, yeast strain MD73 (Lrs4-3HA, Brn1-13x-myc) was grown to OD_600_ of 0.6-0.8, washed, and frozen at −80°C. Cell pellets were resuspended in lysis buffer (50 mM HEPES-KOH [pH 7.5], 150 mM NaCl, 1 mM EDTA, 10% glycerol, 0.5% Nonidet P-40) containing Complete Protease Inhibitor Cocktail (Roche) and then disrupted with glass beads in a Mini-Beader-16 (Biospec Products). Extracts were centrifuged at 13,000 rpm in a microfuge for 15 min at 4°C. Supernatants were incubated with anti-HA ((#C29F41) or anti-myc (9E10) antibody at 4°C for 2 hr, followed by 30 μl of pre-washed Protein A agarose beads (GE) for 2hr. Beads were collected by centrifugation at 500xg for 5 min, washed twice with wash buffer (50 mM HEPES-KOH [pH 7.5], 150 mM NaCl, 1 mM EDTA), boiled in 1X Laemmli buffer at 95°C for 5 min and run on SDS-PAGE gel for immunoblotting.

### Chromatin Immunoprecipitation

ChIP assays were performed as previously described (Li *et al.*, 2013). Recovered DNA was analyzed by real-time PCR using SensiMix SYBR Hi-ROX kit (Bioline, # QT605-05) on a Step One plus real-time PCR machine (Applied Biosystems). PCR primer sequences used for the study are listed in Table S3. Enrichment was calculated at the respective loci and plotted as percent input. For ChIP-seq experiments, libraries were prepared from the recovered DNA using an Illumina Trueseq ChIP Sample Prep kit (#IP-202-1024) and TrueSeq standard protocol. Single-end sequencing was completed on an Illumina NextSeq 500 at the UVA Genomic Analysis and Technology Core, RRID:SCR_018883. Biological duplicate fastq files for each sample were merged and reads were aligned to the sacCer3 reference genome (release R64-2-1) using Bowtie2 with the following options:—best,—stratum,—nomaqround, and—m10 (Langmead et al., 2009). The aligned reads were filtered and indexed using SAMtools to generate bam files, which were then converted to bed files using BEDTools (Quinlan and Hall, 2010). The raw and processed datasets used in this study have been submitted to the NCBI GEO database. The ML1 input sample is available at GSE92714. MACS2 was used to call peaks with the following options:—broad,—keep-dup, -tz 150, and -m 3, 1000 (Zhang et al., 2008). Ctf3 peaks in the WT backgrounds were subtracted from the WT Brn1-13xMyc, WT Lrs4-13xMyc and *lrs4*Δ Brn1-13xMyc peaks, respectively, using BEDTools “intersect” with the–v option. DiffBind was used to call for peaks that were differentially bound in WT Brn1-13xMyc and *lrs4*Δ Brn1-13xMyc samples.

### ChIP-reChIP

ChIP-reChIP was performed as previously described (Furlan-Magaril et al., 2009). The steps were similar to standard ChIP with the first primary antibody until elution of the chromatin DNA from beads, which was done with 75 μl reChIP elution buffer (Tris-EDTA, 1% SDS, 10 mM DTT, protease inhibitor cocktail) by incubating at 37°C for 30 min. The eluent was transferred to a new microfuge tube and diluted 20 times with dilution buffer (20 mM Tris HCl, pH 8, 150 mM NaCl, 1% Triton X-100, 2 mM EDTA, protease inhibitor cocktail). Another primary antibody was incubated, followed by incubation with Protein G magnetic beads (Thermo Fischer Scientific). Following washes and recovery of DNA with final elution buffer (1% SDS and 100 mM NaHCO_3_), the purified DNA was subjected to qPCR for detecting relative enrichment of co-bound proteins at the indicated loci.

### Micro-C XL

Micro-C XL was performed as previously described (Hsieh *et al.*, 2016), with modifications. For the WT, *lrs4*Δ and *fob1*Δ mutants, 100 ml YPD cultures were grown to mid-log phase. For the Brn1-AID depletion experiment, log phase SC + 2% raffinose cultures were treated or not treated with 0.5 mM auxin for 30 min, followed by the addition of 2% galactose for 1 hr, and then 2% glucose for 30 min to induce switching. All cultures were then crosslinked with 3% formaldehyde for 10 min at 30°C and quenched with glycine. Cells were centrifuged, washed with sterile water, resuspended in 10 ml Buffer Z (1M sorbitol, 50mM Tris, pH 7.4) and 10 mM β-Mercaptoethanol, then spheroplasted by adding 250 μl of 20T 10 mg/ml Zymolyase for 1 hr at 30°C. Cell pellets were washed in 1X PBS and crosslinked with 3 mM EGS (ethylene glycol bis(succinimidyl succinate)) for 40 min at 30°C, quenched by addition of glycine, washed, and stored at −80°C. The cell pellets were resuspended in 50 μl of MBuffer1 and digested with an appropriate amount of MNase (Worthington, #LS004798) to obtain >95% mononucleosomes. The chromatin pellet from four combined reactions was then used for Micro-C XL following the published protocol (Hsieh *et al.*, 2016). Libraries of the recovered ligated dinucleosomal DNA were prepared using NEBNext Ultra II kit (#E7645S, New England Biolabs). Paired-end sequencing was performed on an Illumina NextSeq 500 at the UVA Genome Analysis and Technology Core, RRID:SCR_018883.

Micro-C XL raw data was filtered and mapped into binned and iteratively corrected genome matrix files using the HiC-Pro pipeline (Servant et al., 2015) on UVA’s high-performance computing Rivanna cluster. Matrix file outputs were further analyzed with the HiTC package in R studio to create the genome and individual chromosome interaction maps presented in this work (Servant et al., 2012). A custom awk script was used to parse out the top 10% of annotated genomic interactions presented in Figures S3A and B and is available upon request. BART3D was used to calculate Differential Chromatin Interaction (DCI scores).

### Switching assay and donor preference assay

The donor preference assay was performed as previously described (Li et al., 2012). The mating-type switching time course with strains carrying pGAL-HO-URA3 was performed by isolating genomic DNA from cells harvested at each timepoint and the switching detected by relative abundance of *MAT*a and *MAT*α cassettes detected by PCR (Li *et al.*, 2019).

### RNA extraction and RT-qPCR

Total RNA was extracted using the hot acid-phenol method. 2 µg of RNA was then reverse transcribed with oligo(dT) using a Verso cDNA synthesis kit (Thermo Fisher Scientific). The qPCR reactions were performed using a SensiMix SYBR Hi-ROX kit (Bioline, QT605-05) on a StepOnePlus Real-Time PCR System (Applied Biosystems). The qPCR fold induction was calculated using the ΔΔCt method relative to the loading control *ACT1* and then normalized to 1.0 for a specific condition in each experiment. Primers are listed in Table S3.

### GST-Fob1p purification

Yeast strains were cultured in 2L SC-ura with 2% raffinose to log phase, then centrifuged at 3500 rpm in a F9-4×1000Y rotor (Piramoon Technologies) for 15 min at room temperature. Cells were resuspended and incubated in SC-ura with 2% galactose to induce GST-Fob1 expression for 4 hrs. Following centrifugation and washing, cell pellets were resuspended in 20 ml lysis buffer (50 mM HEPES, pH 7.6, 10% glycerol, 10 mM EDTA, 0.5M NaCl, 1% Triton X-100, 5 mM DTT, 1X protease inhibitor cocktail, 1 mM PMSF) and disrupted with glass beads using a BeadBeater (Biospec Products). The lysate was centrifuged at 15000 rpm in a Sorvall SS-34 rotor for 10 min at 4°C and the soluble extracts diluted with lysis buffer to a final NaCl concentration of 0.1M, then incubated with 2 ml glutathione 4B-sepharose (GE, 17-0756-01) for 4 hr at 4°C. The resin was washed 2 times with wash buffer (50 mM HEPES, pH 7.6, 10% glycerol, 10 mM EDTA, 0.1M NaCl, 1% Triton X-100, 5 mM DTT, 1X protease inhibitor, 1 mM PMSF) and the GST fusion protein was eluted with 400 µl elution buffer (50 mM HEPES, pH 7.6, 10% glycerol, 100 mM NaCl, 1mM DTT, 10 mM glutathione). The eluate was concentrated in a 3K MWCO filter Amicon Ultra centrifugal filter at 14000 g for 10 min at 4°C. Isolation of the fusion protein was confirmed by immunoblot with α-Fob1 antibody (Santa Cruz, sc-98575, 1:1000 dilution).

### EMSA

DNA fragments for EMSA were PCR amplified from genomic DNA with 5’-biotinylated primers (Table S3), then purified from a 12% polyacrylamide gel. EMSA was performed with a LightShift Chemiluminescent EMSA kit (Thermo Fischer, 20148). 5 ng of probe was incubated with 60 ng GST or GST-Fob1p in 20 µl 1X buffer (10 mM Tris, pH 7.5, 1 mM EDTA, 0.1M KCl, 0.1mM DTT, 5% glycerol) and incubated at 30°C for 30 min. As competitor, 50 ng or 200 ng unlabeled probe was added. The RFB fragment was used a positive control for direct interaction (Kobayashi, 2003), while a nuclear extract supplied with the kit was used as a positive gel shift control. Samples were run on a 6% polyacrylamide gel in 0.5X TBE, transferred to Immobilon-Ny^+^ at 380mA (∼100V) for 30 minutes using mini Trans-blot electrophoretic system (Bio-Rad) and autocrosslinked at 120 mJ/cm^2^ in a Stratagene Stratalinker 1800. The biotin-labeled DNA was detected by using LightShift Chemiluminescent EMSA reagents according to the manufacturer’s instructions.

## Acknowledgments

We thank Jonathan Dinman, Danesh Moazed, Vincent Guacci, Jessica Tyler, James Haber, Alan Hinnebusch, and Kevin Lynch for kindly providing yeast strains and plasmids. We also thank Lindsey Power for critical reading of the manuscript, and other Smith lab members for helpful discussions. This work was supported by NIH grants 5R01GM127394 and 2R01GM075240 to J.S.S. and R35GM133712 to C.Z. The research was also supported by NCI Cancer Center Support Grant 5P30CA044579 and by the UVA Genome Analysis and Technology Core, RRID:SCR_018883.

## Author contributions

M.D., M.L., and J.S.S. conceptualized the study. M.D. and J.S.S. wrote the manuscript. M.D., R.D.F., S.S., Z.W., and M.L. performed the methodology. M.D., R.D.F., and J.S.S. were responsible for the resources. M.D., R.D.F., and Z.W. curated and visualized the data. J.S.S. and C.Z. supervised the study. J.S.S. and C.Z. acquired the funding.

## Supplemental Information

1. Supplemental Table S1. List of differential Brn1-myc ChIP-seq peaks in *lrs4*Δ strain compared to WT. <u>Excel spreadsheet</u>

**Supplemental Table S2.**
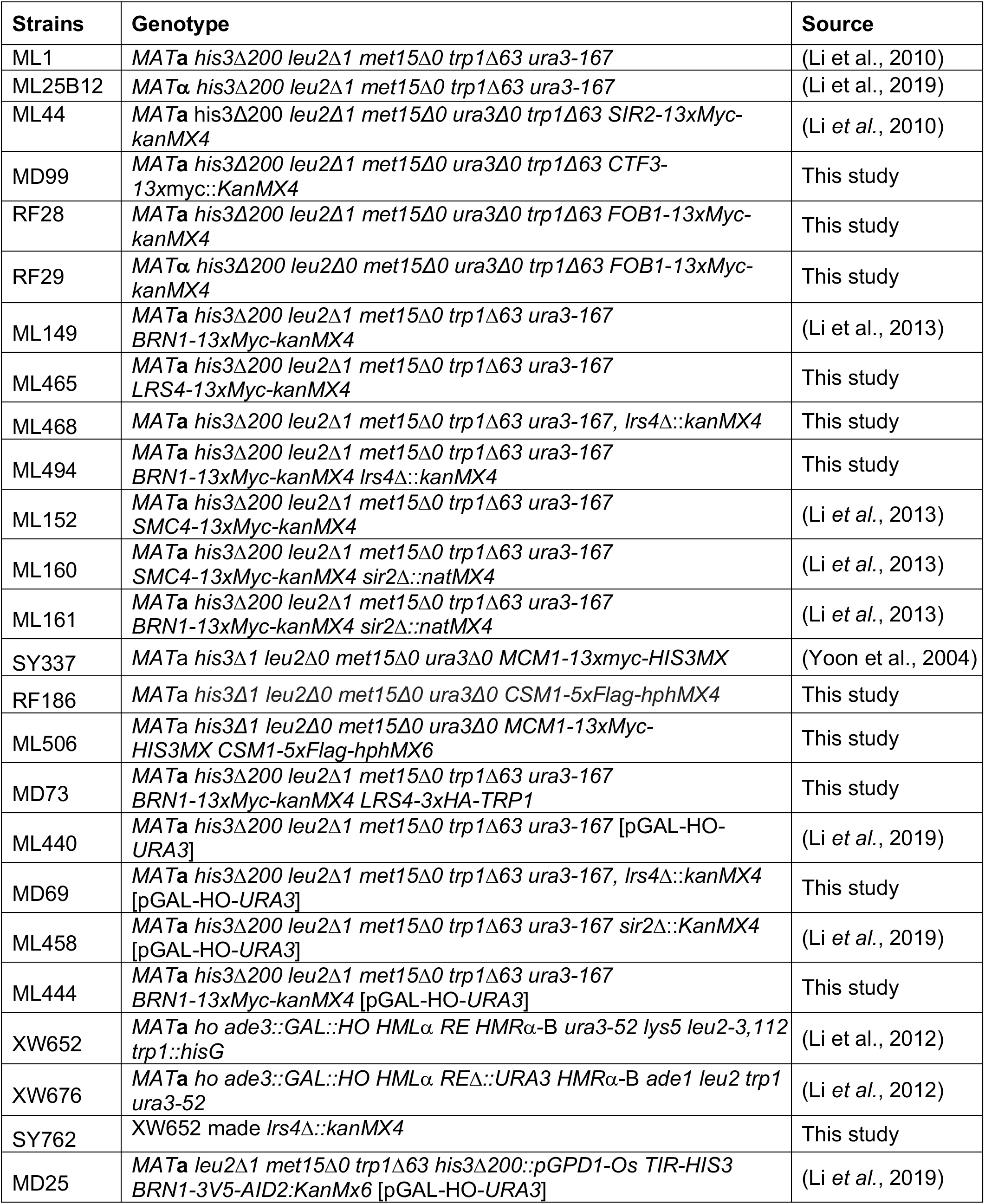
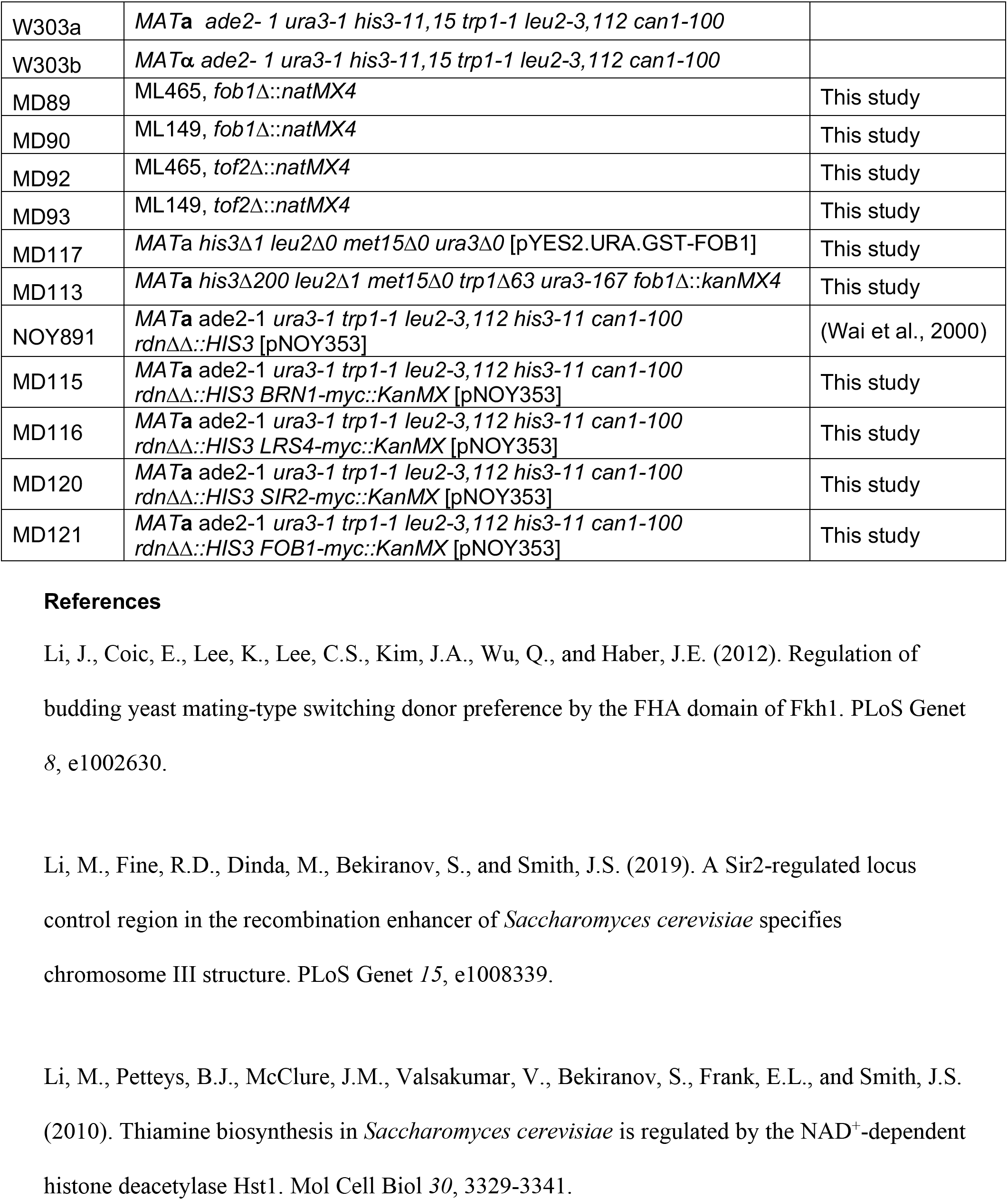
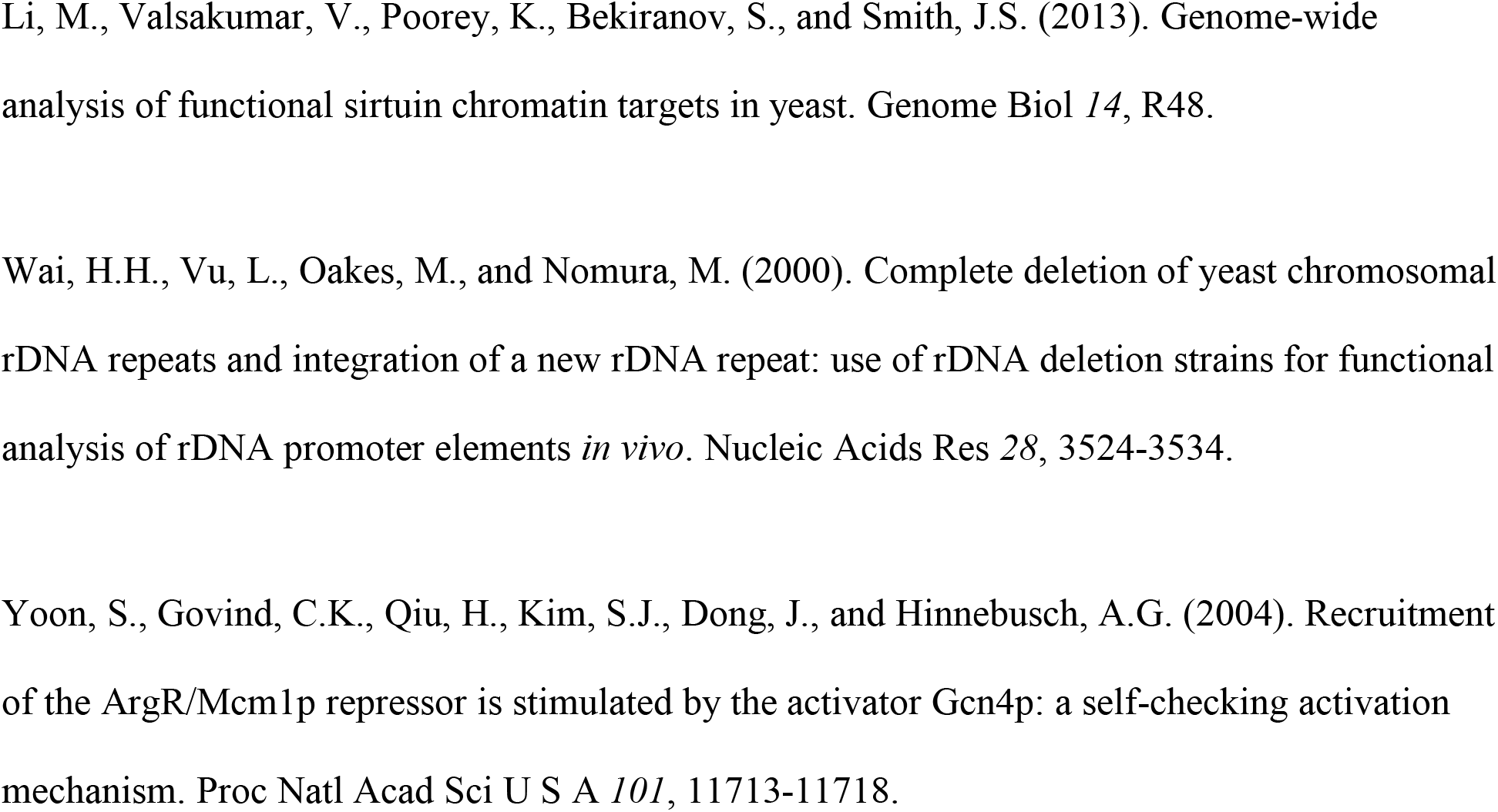
Yeast Strains

**Supplemental Table S3.**
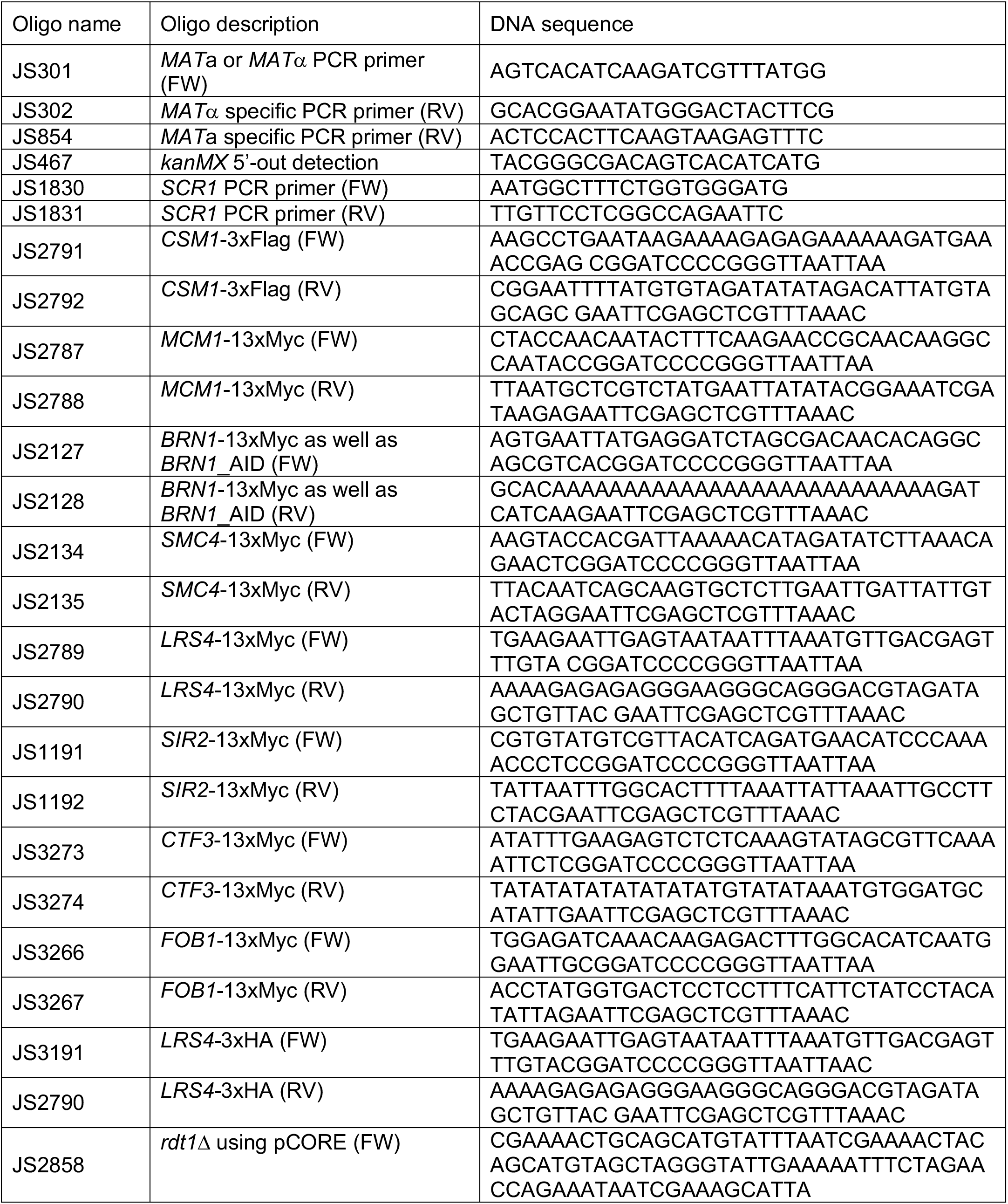
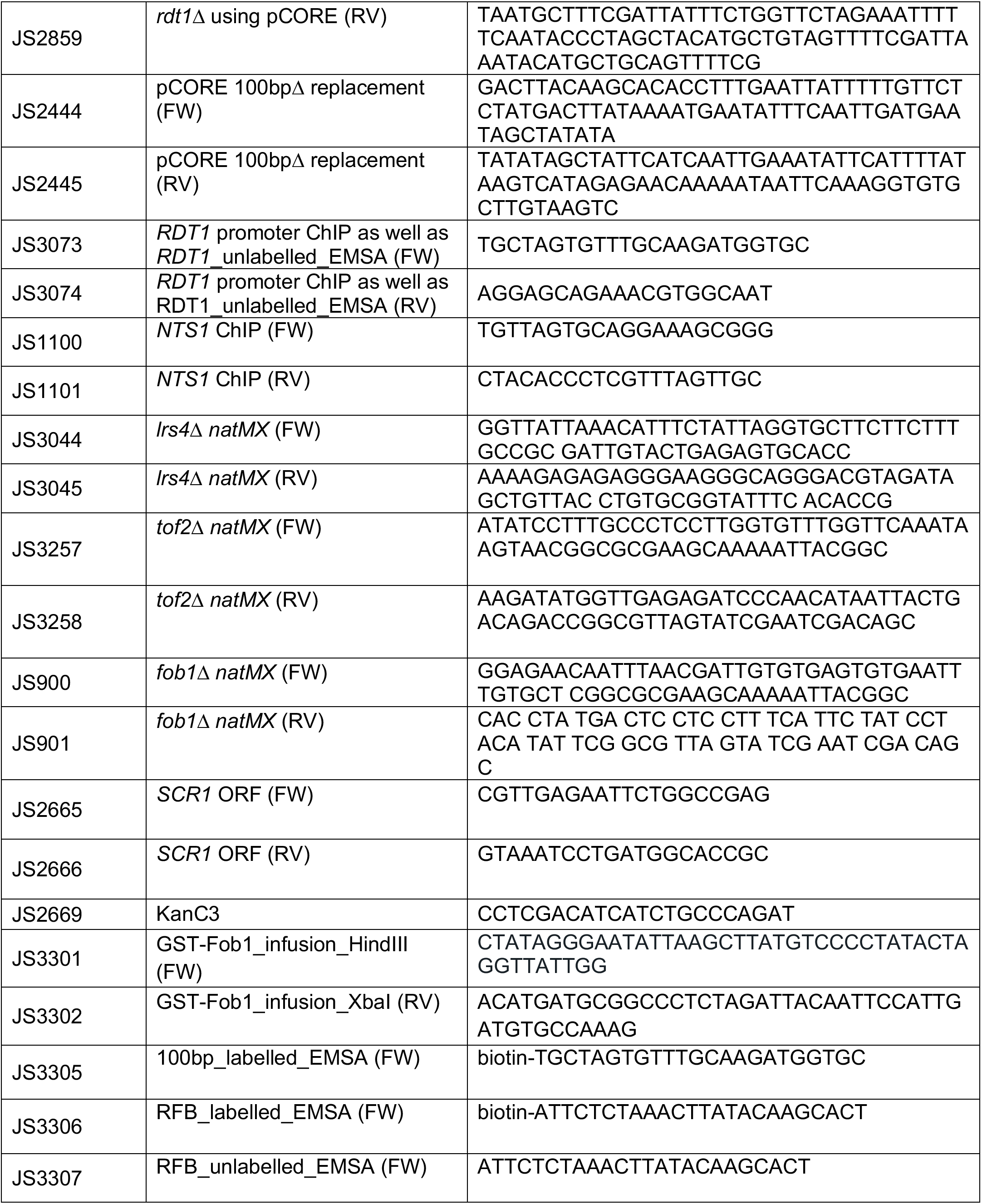
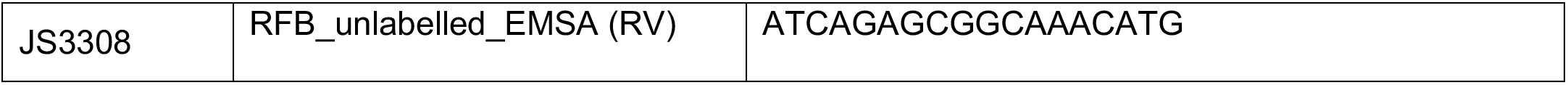
Oligonucleotides

**Figure S1.**
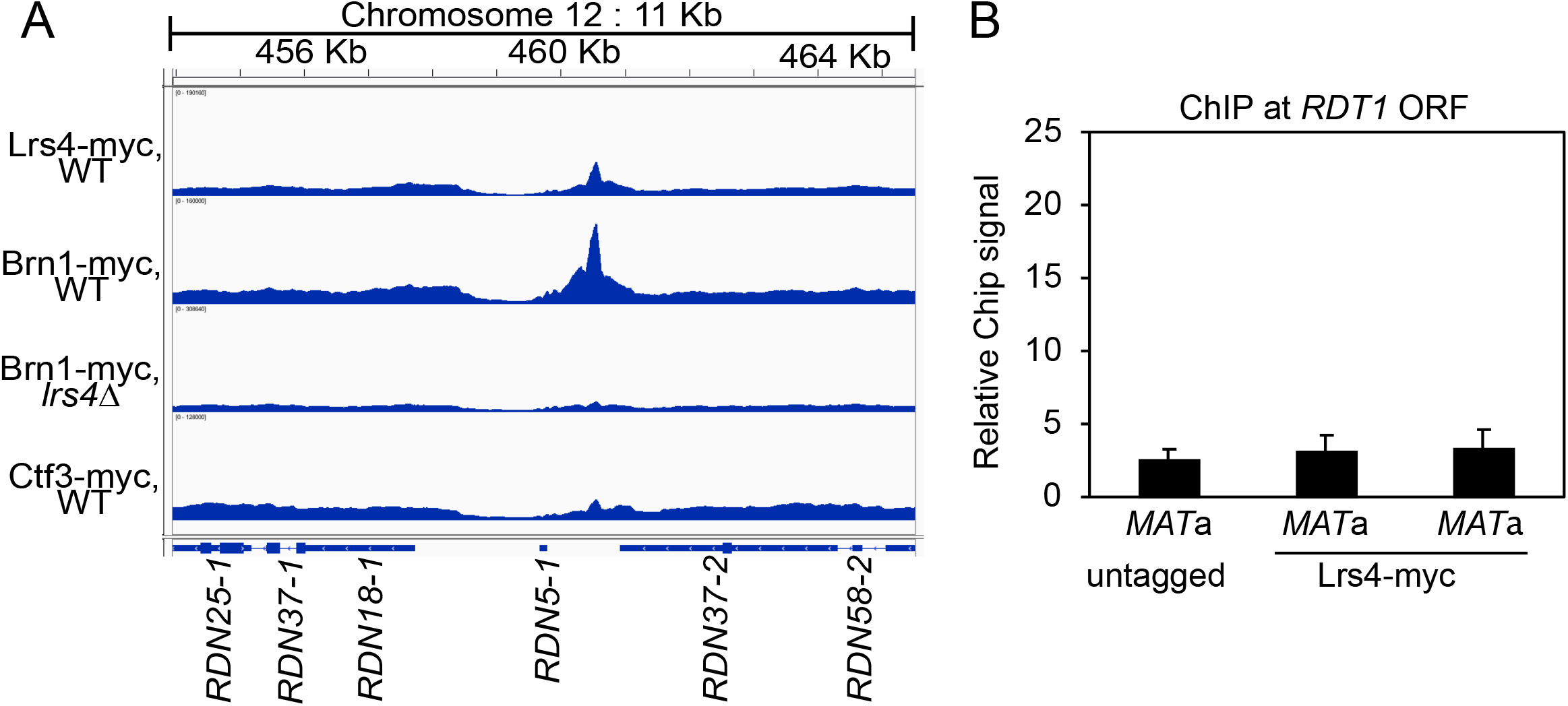
ChIP-seq enrichment peaks for Lrs4-myc and Brn1-myc at the rDNA locus. (*A*) IGV snapshot for chromosome XII of ChIP-seq data showing the enrichment of Brn1-myc and Lrs4-myc, and loss of Brn1-myc binding in the *lrs4*Δ mutant. Ctf3-myc is used as a control for non-specific binding independent of centromeres. Tracks were normalized to total read count and comprise the average enrichment across all rDNA repeats. (*B*) ChIP assay with Lrs4-myc showing lack of enrichment on the *RDT1* open reading frame in *MAT*a and *MAT*α cells.

**Figure S2.**
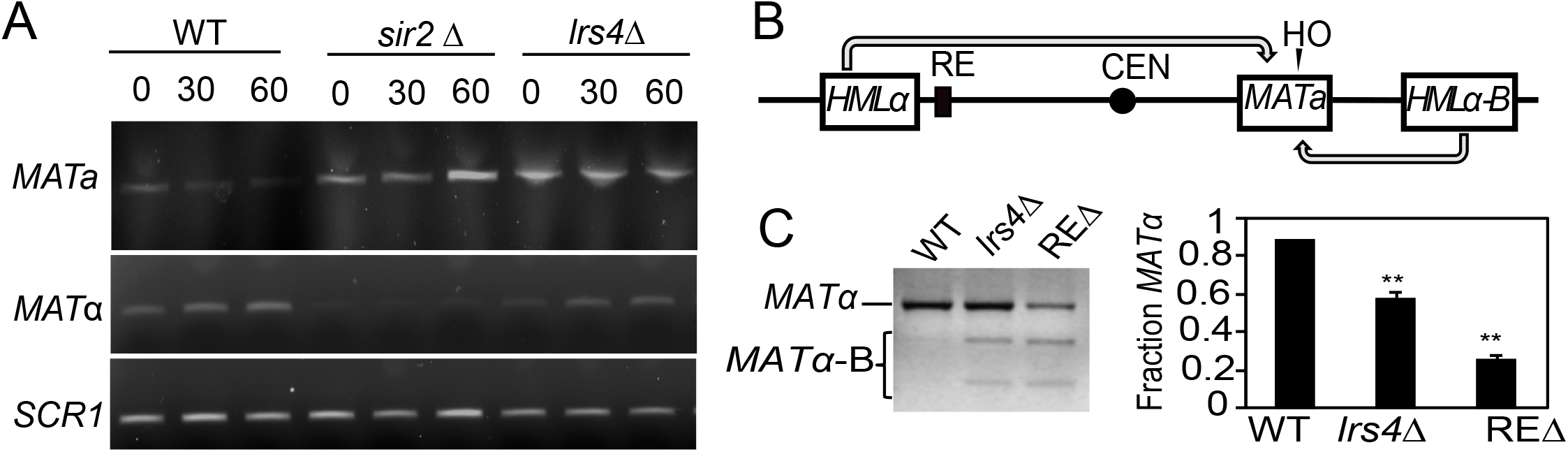
Regulation of mating-type switching by Lrs4. (*A*) Time-course of mating-type switching in WT (ML447), *sir2*Δ (ML458) and *lrs4*Δ (MD69) showing PCR detection of *MAT*a and *MAT*α. *SCR1* is used as a control for input genomic DNA. (*B*) Schematic of the donor preference assay that distinguishes usage of *HML*α or an engineered *HMR*α-B cassette as the donor template by digestion of a *MAT*α PCR product with *Bam*HI. (*C*) Donor preference assay with WT (XW652), *lrs4*Δ (SY762), or *RE*Δ (XW676) showing *Bam*HI digestion of the *MAT*α PCR product after switching. Upper band indicates normal donor preference switching using *HML*α as the template. Digested bands indicate improper switching using *HMR*α-B as the template. The fraction of normal upper band *MAT*α PCR products is quantified in the bar graph. Significant decreases = **p<0.005.

**Figure S3.**
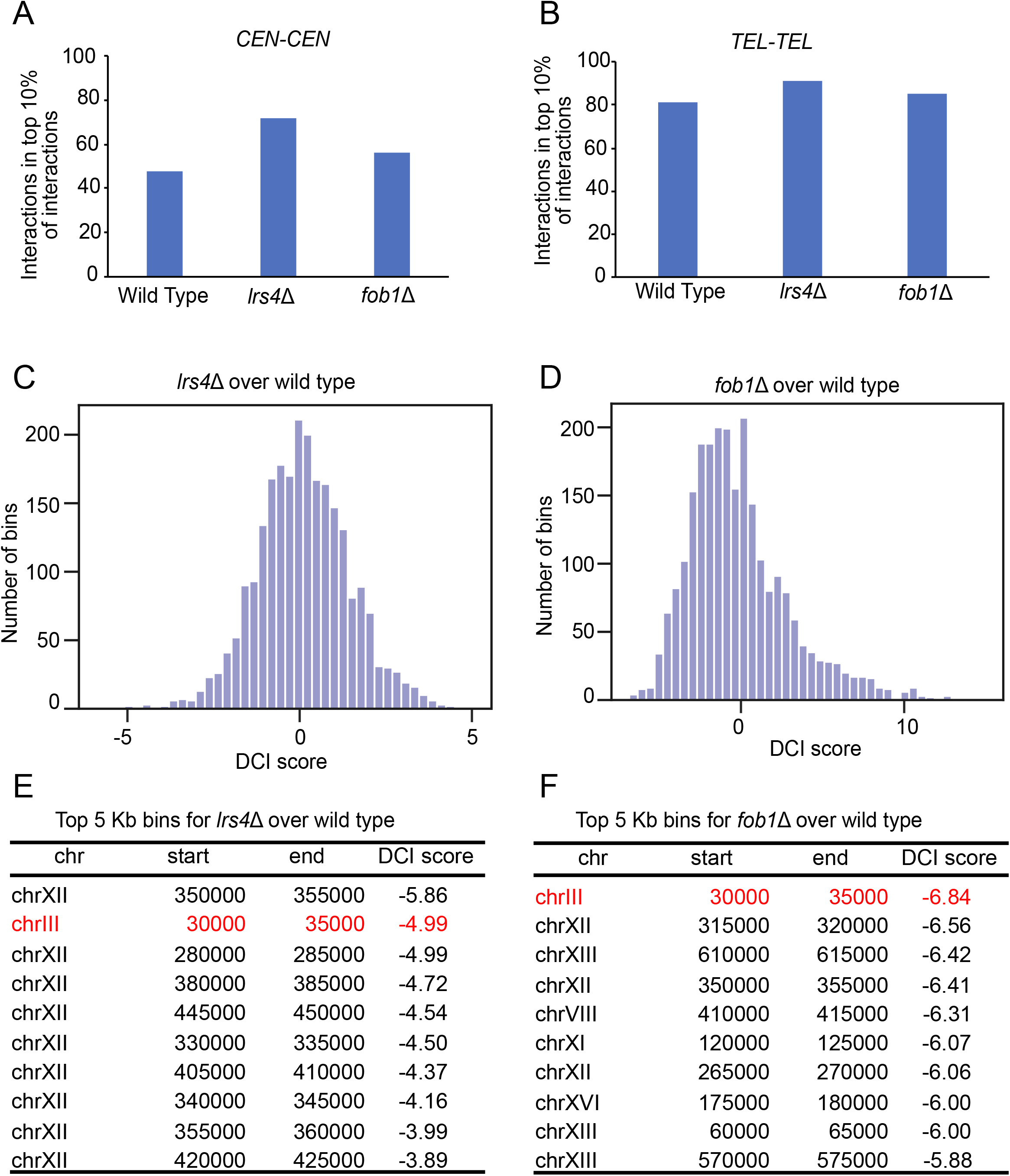
Differential Chromatin Interaction (DCI) analysis of *lrs4*Δ and *fob1*Δ mutants. (*A-B*) Genomic interactions between centromeres (*A*) and telomeres (*B*) across 10 kb bins in *lrs4*Δ and *fob1*Δ. p < 0.05 calculated using a difference of means permutation test of *CEN/TEL* bins versus the rest of the genome. (*C-D*) Distribution of DCI scores for all 5 kb bins in the genome for *lrs4*Δ (*C*) and *fob1*Δ (*D*) compared to Wild Type. (*E-F)* Ranked list of 5 kb bins with the strongest negative DCI scores. The 5kb bin containing *RDT1* highlighted in red.

